# The Maintenance of Muscle Mass Is Independent of Testosterone in Adult Male Mice

**DOI:** 10.1101/2020.09.24.311266

**Authors:** Arik Davidyan, Keith Baar, Sue C. Bodine

## Abstract

Testosterone is considered a potent anabolic agent in skeletal muscle with a well-established role in adolescent growth and development in males. However, alterations in the role of testosterone in the regulation of skeletal muscle mass and function throughout the lifespan has yet to be established. While some studies suggest that testosterone is important for the maintenance of skeletal muscle mass, an understanding of the role this hormone plays in young, adult, and old males with normal and low serum testosterone levels is lacking. We investigated the role testosterone plays in the maintenance of muscle mass by examining the effect of orchiectomy-induced testosterone depletion in C57Bl6 male mice at ages ranging from early postnatal through old age; the age groups we used included 1.5-, 5-, 12-, and 24-month old mice. Following 28 days of testosterone depletion, we assessed mass and fiber cross-sectional-area (CSA) of the tibialis anterior, gastrocnemius, and quadriceps muscles. In addition, we measured global rates of protein synthesis and degradation using the SuNSET method, western blots, and enzyme activity assays. 28 days of testosterone depletion resulted in smaller muscle mass in the two youngest cohorts but had no effect in the two older ones. Mean CSA decreased only in the youngest cohort and only in the tibialis anterior muscle. Testosterone depletion resulted in a general increase in proteasome activity at all ages. We did not detect changes in protein synthesis at the terminal time point. This data suggest that within physiological serum concentrations, testosterone is not important for the maintenance of muscle mass in mature male mice; however, in young mice testosterone is crucial for normal growth.

## Introduction

Skeletal muscle is a critical tissue for movement and posture, as well as, other bodily functions that impact quality of life and independence including serving as a protein reservoir, aiding with blood glucose homeostasis, contributing to heat generation, and regulating metabolism (1). Skeletal muscle mass is often used as a marker of skeletal muscle health throughout life (2); however, the cumulative processes and mechanisms that are important for skeletal muscle mass accretion and maintenance are incompletely understood. Moreover, we do not fully understand the mechanisms responsible for the loss of skeletal muscle mass and function with age.

Skeletal muscle mass is governed by the delicate balance between myofibrillar protein synthesis and degradation (3), and many factors influence this balance including external loading, neural activity, nutrition and hormones. Testosterone is a steroid hormone derived from cholesterol that is linked to a variety of physiological functions including sexual function and development, systemic metabolism, cognitive function, as well as bone and muscle growth and maintenance (4,5). Because of its presumed role in muscle growth, there is much interest in the potential role testosterone might have in maintenance of muscle mass in adults, adaptive growth in response to anabolic signals such as increased load, recovery of muscle mass following injury, and prevention of muscle mass loss during disuse and aging (6,7). Testosterone is thought to affect both myofibrillar protein degradation and protein synthesis pathways (8). Currently, testosterone is hypothesized to promote protein synthesis via activation of Akt and mTORC1 (9–11), as well as inhibition of protein degradation through the ubiquitin-proteasome-system (10,12,13) through its antagonistic actions of the glucocorticoid receptor (14–16). However, the specific mechanisms that mediate the effects of testosterone on skeletal muscle are still unclear. In skeletal muscle, testosterone is thought to be highly anabolic. However, the degree to which testosterone exerts its influence on skeletal muscle mass throughout the lifespan is unclear. Many studies that investigate the effects of testosterone depletion use either young growing animals (11,17,18) or old animals (9). Studies that examine the actions of testosterone on skeletal muscle at more than two life stages are uncommon. Those studies that examine the effect of testosterone in either young or old animals often contradict one another and may not accurately reflect the role of testosterone in fully mature adult animals. Two weeks of testosterone depletion by orchiectomy in young (8-week old) C57Bl6/6J male mice as well as young adult (10-month old) Fischer 344 male rats has been shown to result in ∼50% smaller muscle mass (11,19). In contrast, another study in young male Sprague-Dawley rats (8-week old) showed no change in lean body mass following 2 and 4 weeks of castration (20). Moreover, 16 weeks following castration in 20 weeks old mice demonstrated no difference in lean mass (21). Clearly, there is a need to understand how testosterone contributes to skeletal muscle health and function throughout the lifespan, i.e., juvenile, adult, and aging animals. Therefore, the primary objective of this study was to investigate the role testosterone plays in the regulation of muscle mass by examining the effect of orchiectomy-induced testosterone depletion in male mice at ages ranging from early postnatal to old age. We hypothesized that phenotypic outcomes in skeletal muscle following testosterone depletion would differ between male mice of different ages and developmental stages. Moreover, we hypothesized that any potential changes in skeletal muscle mass following castration would be dictated by alterations to protein degradation.

## Materials and Methods

### Animals and ethical approval

Male C57Bl6 mice were obtained from Charles River Laboratories at four different age groups (1-, 3.5-, 4.5-, and 11.5-months old) and from the National Institute of Health (NIA Rodent Colony) at 23.5-months of age. All animals were allowed two weeks of acclimatization prior to the start of experimentation. Three to four animals were housed together in ventilated cages at a temperature of 25°C and fed ad libitum. Animals were subjected to a 12-hour light and 12-hour dark cycle. Animal procedures were approved by the Institutional Animal Care and Use Committee at the University of California, Davis.

### Castration

To deplete testosterone, animals from each age group were randomized to either a sham castration group (Sham) or Castration (Cast), resulting in the following eight groups: 1.5-months old Sham and Cast (n=8 and n=12 respectfully), 5-months old Sham and Cast (n=6 for both), 12-months old Sham and Cast (n=7 for both), and 24-months old Sham and Cast (n=8 and n=4 respectfully). In addition, a 4-month old control group was subjected to sham castration. At day 0, animals were sedated with 2–3% inhaled isoflurane and placed in a sterile surgical field. Using aseptic surgical procedures, an incision (∼2cm) was made to the skin in the lower abdomen followed by blunt dissection and an incision to the lower abdominal muscles. The left testicle was pulled out of the abdominal cavity and was removed following an occlusion of the spermatic cord. The teste-less spermatic cord was returned into the abdominal cavity and the procedure was repeated on the right side. The abdominal muscle was closed with an interrupted suture and the cutaneous incision was closed with a continuous subcuticular suture. Mice were given an analgesic (buprenorphine, 0.1mg/kg) immediately following the surgery and returned to their cages. Animals were returned to their cage and fed ad libitum for 28 days prior to euthanasia. Sham operation were identical, excluding the occlusion of the spermatic cord and the removal of the testes. If needed, additional analgesic was given for 48 hours following the surgery.

### Tissue Collection

Following completion of the treatment period, mice were anesthetized with 2.5% inhaled isoflurane. Under anesthesia, hindlimb muscles were excised bilaterally, along with the right perirenal fat pad. All tissues were weighed and frozen in liquid nitrogen for biochemical analyses. Muscles used for histological analysis were pinned on cork at a length approximating Lo (resting length) and frozen in liquid nitrogen-cooled isopentane. Mice were euthanized following tissue collection by exsanguination.

### Serum T measurement

Blood samples were centrifuged at 2,000 RPM for 10 minutes at 4°C. Supernatant was collected and frozen. Samples were analyzed for total serum testosterone levels using a mouse/rat testosterone ELISA kit from Calbiotech Inc. (Spring Valley, CA # Cat. # TE187S-100) as indicated by the company. Samples were analyzed in duplicates.

### Rate of Protein Synthesis Measurements

Protein synthesis was measured in mice using the SUnSET method as previously described (22). Exactly 30 min before the muscles were excised, mice were given an intraperitoneal injection of 0.04μmol puromycin per gram of body weight dissolved in 100μl of sterile phosphate buffered saline (PBS). The amount of incorporated puromycin was analyzed by western blot as described below.

### Immunoblotting

Frozen muscles were powdered and homogenized in sucrose lysis buffer (50mM Tris at pH 7.5, 250mM sucrose, 1 mM EDTA, 1mM EGTA, 1% Triton, 50mM NaF, 5mM Na2(PO4)2, and protease inhibitor). The supernatant was collected following centrifugation at 8,000g for 10 min and protein concentrations were determined in triplicate using the Bradford method (Bio-Rad). Ten micrograms of protein were subjected to SDS-PAGE on 4–20% Criterion TGX Stain-Free Protein Gel (Bio-Rad) and transferred to polyvinylidene diflouride (PVDF) membrane, which was previously activated for 10 minutes with 100% ethanol. Membranes were blocked in 1% skim milk dissolved in Tris-buffered saline with 0.1% Tween-20 (TBST) for one hour and then probed with primary antibodies overnight at 4°C. The following day, membranes were washed and incubated with HRP conjugated secondary antibodies at 1:10,000 for one hour at room temperature. Immobilon Western Chemiluminescent HRP substrate (Millipore) was then applied to the membranes for protein band visualization by chemiluminescence. Image acquisition and band quantification was performed using the ChemiDoc MP System and Image Lab 5.0 software (Bio-Rad). Total protein staining (whole lane) of the membrane was used as the normalization control for all blots. The following primary antibodies were used in this study at a concentration of 1:1000: Puromycin (Millipore, catalog #MABE343) and Phospho-4E-BP1 (Cell Signaling, #2855).

### Proteasome Activity

20S and 26S β5 proteasome activity were measured as described previously (23). Briefly, proteasome activity was measured in the supernatant after 30 min centrifugation at 12,000 g following homogenization in 300μl of buffer containing 50mM Tris, 150mMNaCl, 5mMMgCl2, 1mMEDTA, and 0.5mMDTT at pH 7.5. The chymotrypsin (β5)-like activities were assayed using 10μg of protein and the fluorescently tagged substrate SUC-LLVY-AMC (Bachem). Both assays were carried out in a total volume of 100μl. The 26S ATP-dependent assay was performed in homogenization buffer with the addition of 100μM ATP. The 20S ATP-independent assay was carried out in assay buffer containing 25mM HEPES, 0.5mM EDTA, and 0.001% SDS (pH 7.5). Each assay was conducted in the absence or presence of the proteasome inhibitor Bortezomib at a final concentration of 2mM. The activity of the 20S and 26S proteasome was measured by calculating the difference between fluorescence units recorded with or without the inhibitor in the reaction medium. Released AMC was measured using a Fluoroscan Ascent fluorometer (Thermo Electron) at an excitation wavelength of 390nm and an emission wavelength of 460 nm. Fluorescence was measured at 15-min intervals for 75 minutes and the last time point was used for analysis. All assays were linear in this range and each sample was assayed in triplicate and all samples for a single analysis were on the same plate.

### Immunohistochemistry

Serial cross sections (10 μm) were cut from the TA and GA muscles using a Leica CM 3050S cryostat (Leica Microsystems). Muscle sections were fixed in cold acetone for 5 minutes at −20°C, followed by 3- and 5-minute phosphate-buffered saline washes. Sections were then incubated with Alexa Fluor® 488 conjugated goat anti-rat immunoglobulin G (IgG; [H+L]; 1:100, Life Technologies) for 1 hour at room temperature. After 3 five-minute phosphate-buffered saline washes, slides were coverslipped using ProLong Gold Antifade reagent with DAPI (Life Technologies). Slides were imaged on a Zeiss Axio Imager.M1 fluorescent microscope using the EC Plan-Neofluar 20× objective. Images were analyzed using FIJI software.

### Fiber Cross-Sectional Area

Serial cross sections (10 μm) were cut from the plantaris using a Leica CM 3050S cryostat. To determine fiber-type-specific cross-sectional area (CSA), plantaris muscle sections were fixed in cold acetone for 5 min at −20°C, followed by three 5-min washes with phosphate-buffered saline with 0.1% Tween 20. Sections were blocked in 5% normal goat serum in phosphate-buffered saline with 0.1% Tween-20 for 30 min at room temperature. After three washes, sections were blocked in Mouse on Mouse blocking buffer made of 2.5ml PBS and 60μl of M.O.M. blocking Reagent (Vector Laboratories, #BMK2202) and then incubated in primary antibody overnight at 4°C; BA-F8 (myosin heavy chain slow type, Mm, IgG2B), SC-71 (myosin heavy chain 2A, Mm, IgG1), and BF-F3 (myosin heavy chain 2B, Mm, immunoglobulin M) were diluted 1:250 in blocking buffer. All anti-myosin heavy chain antibodies were deposited to the Developmental Studies Hybridoma Bank (Iowa City, IA) by Stefano Schiaffino. A polyclonal laminin antibody (1:500, Sigma, #L9393) was included for the determination of CSA. After incubation in primary antibody, sections were incubated in secondary antibody for an hour at room temperature and then coverslipped using ProLong Gold Antifade reagent (Life Technologies, #P36930). For simultaneous detection of multiple mouse primary antibodies, fluorescently conjugated goat-anti-mouse immunoglobulin-specific secondary antibodies were used (Alexa Fluor 350, 488, and 555, Life Technologies). Goat-anti-rabbit AlexaFluor 647 secondary was used to detect laminin. Slides were imaged using a Leica DMi8 inverted microscope using ×10 objective, processed using the Leica LAS X software, and analyzed using the FIJI software. For each animal, all fibers from a cross section were analyzed. For the tibialis anterior muscles, the total number of fibers analyzed was >2000 per muscle. For the gastrocnemius sections the total number of fibers analyzed was >4000 per muscle.

### Statistical Analysis

All data were analyzed using two-way ANOVA using GraphPad Prism software (GraphPad Software, Inc., La Jolla, CA) with Factor 1 being testosterone and Factor 2 being age. Prior to any comparison, Grubbs’ analysis was performed to detect outliers with Alpha = 0.05. Tukey’s post hoc analysis was used to determine differences when interactions existed. Statistical significance (alpha) was set at p<0.05. Statistical significance is presented as following in all Figures; * indicates P< 0.05 for a comparison between age matched experimental and sham groups. ^ indicates P< 0.05 for a comparison between 4-month old and 5-month old groups. Data are presented as mean ± standard error mean (SEM).

## Results

### Effect of testosterone depletion on body and muscle mass throughout lifespan in male mice

Testosterone is thought to contribute to the maintenance of muscle mass in males; therefore, we investigated the effect of 28 days of testosterone depletion on muscle mass in male mice at different ages across the lifespan. Serum testosterone levels peak at 5-months and decrease thereafter (Figure 1B). Orchiectomy decreased serum testosterone levels in all age groups but this was only significant in the 1.5- and 5-month-old cohorts (Figure 1A). Although a two-way ANOVA statistical analysis does not suggest a significant difference between the 12-month-old cohorts, a student T-test as well a multiple T-test analysis suggests that the differences are indeed significant. A significant treatment X age interaction (P=0.004) was observed for body weight (Figure 1A), with testosterone depletion resulting in a significant decrease in the body mass of 12-month-old animals and having no effect in other age groups (Figure 1B). Testosterone depletion significantly decreased perirenal fat pad mass in 12-month-old animals and had no effect in other cohorts (Figure 1C). Mass of the tibialis anterior, gastrocnemius, and quadriceps muscles was lower in 1.5- and 5-month-old mice following one month of testosterone depletion (Figures 1D, 1E, and 1F). Muscle mass was not affected by testosterone depletion in 12- and 24-month-old animals (Figures 1D, 1E, and 1F). The mass of all muscles was significantly lower in 4-month-old sham animals compared to 5-month-old animals sham animals, but was similar in mass to muscles from the 5-month-old castrated animals. The interaction between age and treatment in determining muscle mass was greatest in the tibialis anterior muscle (p=0.011) and weakest in the gastrocnemius muscle (p=0.24).

**Fig 1.**
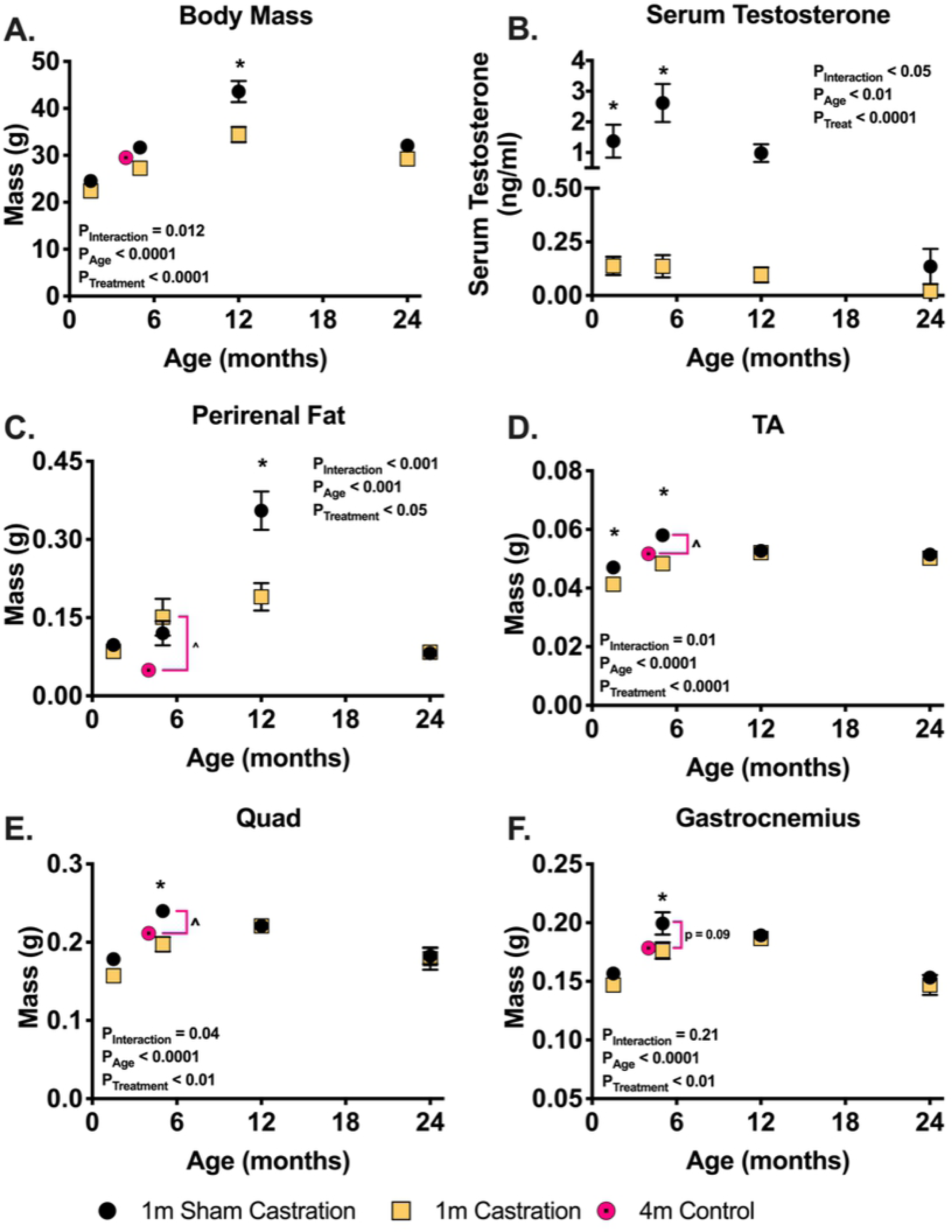
28 days of testosterone depletion selectively decrease muscle mass in young mice and have no effect in older mice. (A) body mass following one month of castration or sham castration surgery. (B) Serum total testosterone measures using an ELISA kit following one month of castration or sham castration surgery. Wet mass of the right perirenal fat pad (C), tibialis anterior (D), gastrocnemius (E), and quadriceps (F) following one month of castration or sham castration surgery. * indicates p < 0.05 between treatment groups of the same age.

### Effect of testosterone depletion on mean and distribution of fiber cross sectional area in male mice

Mean fiber cross sectional area (CSA) of the tibialis anterior was largest in the 5-month-old sham cohort (Figures 2A and 2B). Testosterone depletion significantly decreased the tibialis anterior mean fiber CSA in 1.5-month-old mice but had no significant effect in other age groups (Figures 2A and 2B). The gastrocnemius mean fiber CSA was not affected by treatment. The distribution of fiber cross-sectional areas is shown in Figure 3 for the tibialis anterior and gastrocnemius muscles. The distribution of fiber CSA significantly shifted towards smaller fibers following one month of testosterone depletion in 1.5-month-old animals, an effect that was not seen in any of the other age groups (Figure 3A). In the gastrocnemius muscle, no significant shift in CSA was measured in any age group following 30 days of testosterone depletion (Figure 3B).

**Fig 2.**
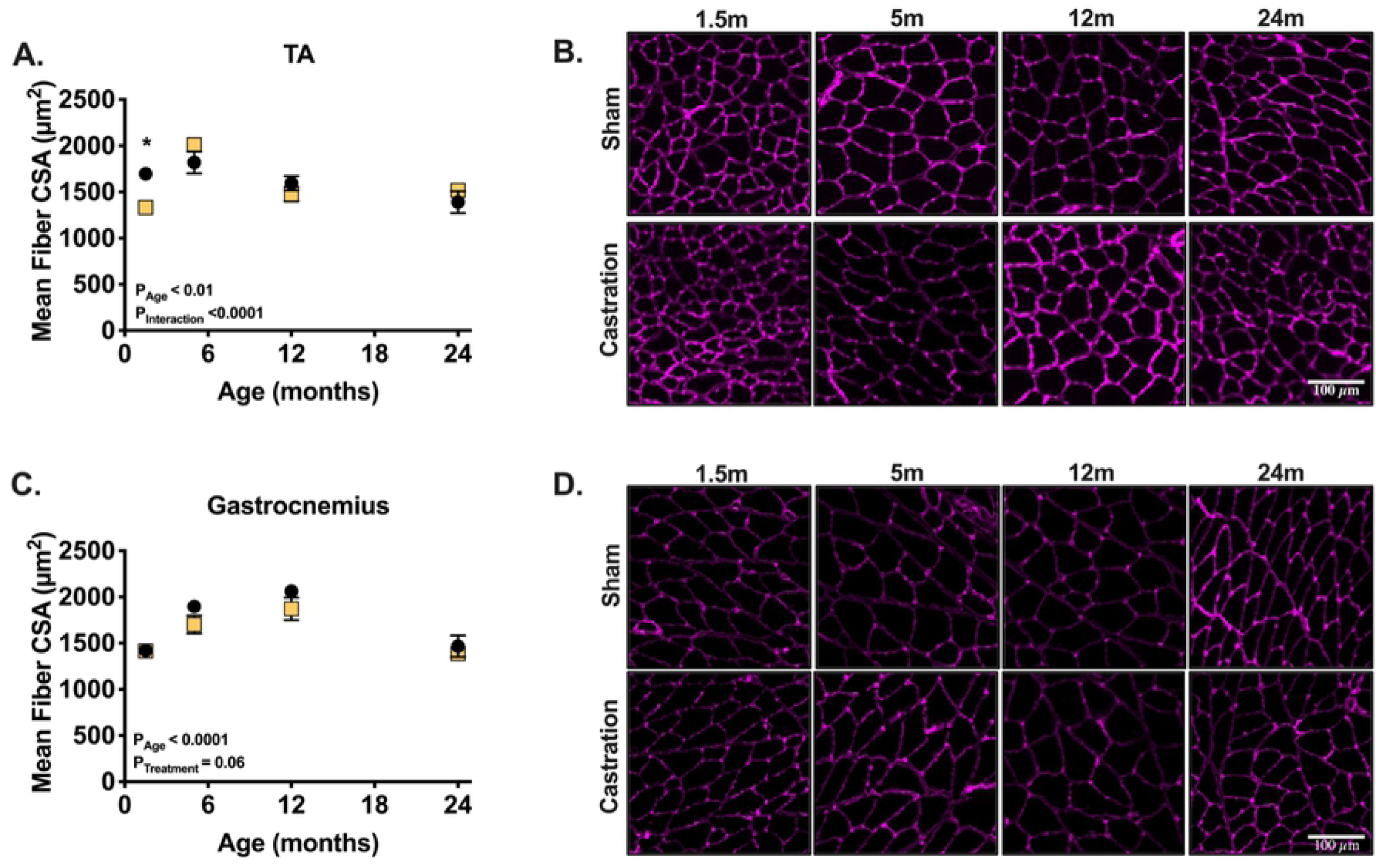
28 days of testosterone depletion decreases fiber CSA only in the TA muscle and only at 1.5-month-old mice. (A) Tibialis anterior CSA following one month of castration or sham castration surgery. (B) Representative images from tibialis anterior histological section stained for laminin. (C) Gastrocnemius CSA following one month of castration or sham castration surgery. (D) Representative images from gastrocnemius histological sections stained for laminin. * indicates p < 0.05 between treatment groups of the same age.

**Fig 3.**
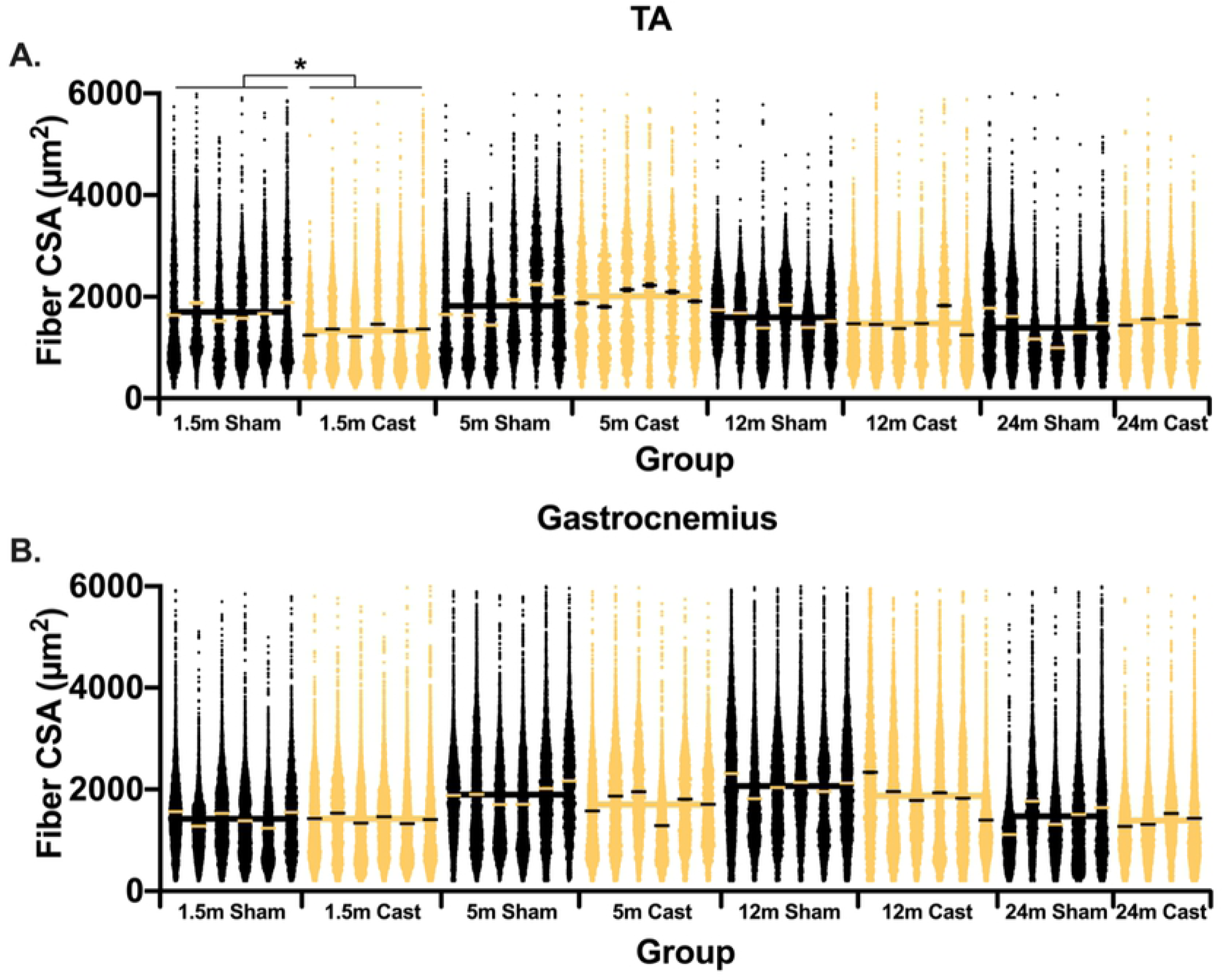
One month of testosterone depletion has no effect on fiber CSA distribution in older mice. (A) Distribution of the tibialis anterior fiber cross section area per animal and group. (B) Distribution of the gastrocnemius fiber cross section area per animal and group. * indicates p < 0.05 between treatment groups of the same age.

### Protein degradation via the ubiquitin-proteasome system following on month of castration in male mice

As protein degradation plays an important role in muscle proteostasis we measured the activity of the 20S and 26S β5 proteasome subunits. The ATP dependent, 26S subunit, decreased with age in the tibialis anterior, gastrocnemius, and quadriceps muscles, with the age-effect being significant in all muscles (Figure 4A, 4B, and 4C). Castration significantly increased the activity of this subunit in the tibialis anterior and the quadriceps muscles. In the gastrocnemius, the 20S, ATP independent, subunit activity decreased as a consequence of age and significantly increased following castration only in the of 5-month-old animals (Figure 4D, 4E, and 4F). Generally, the effects of testosterone depletion across muscles were most visible at 1.5- and 5-months of age (Figure 4).

**Fig 4.**
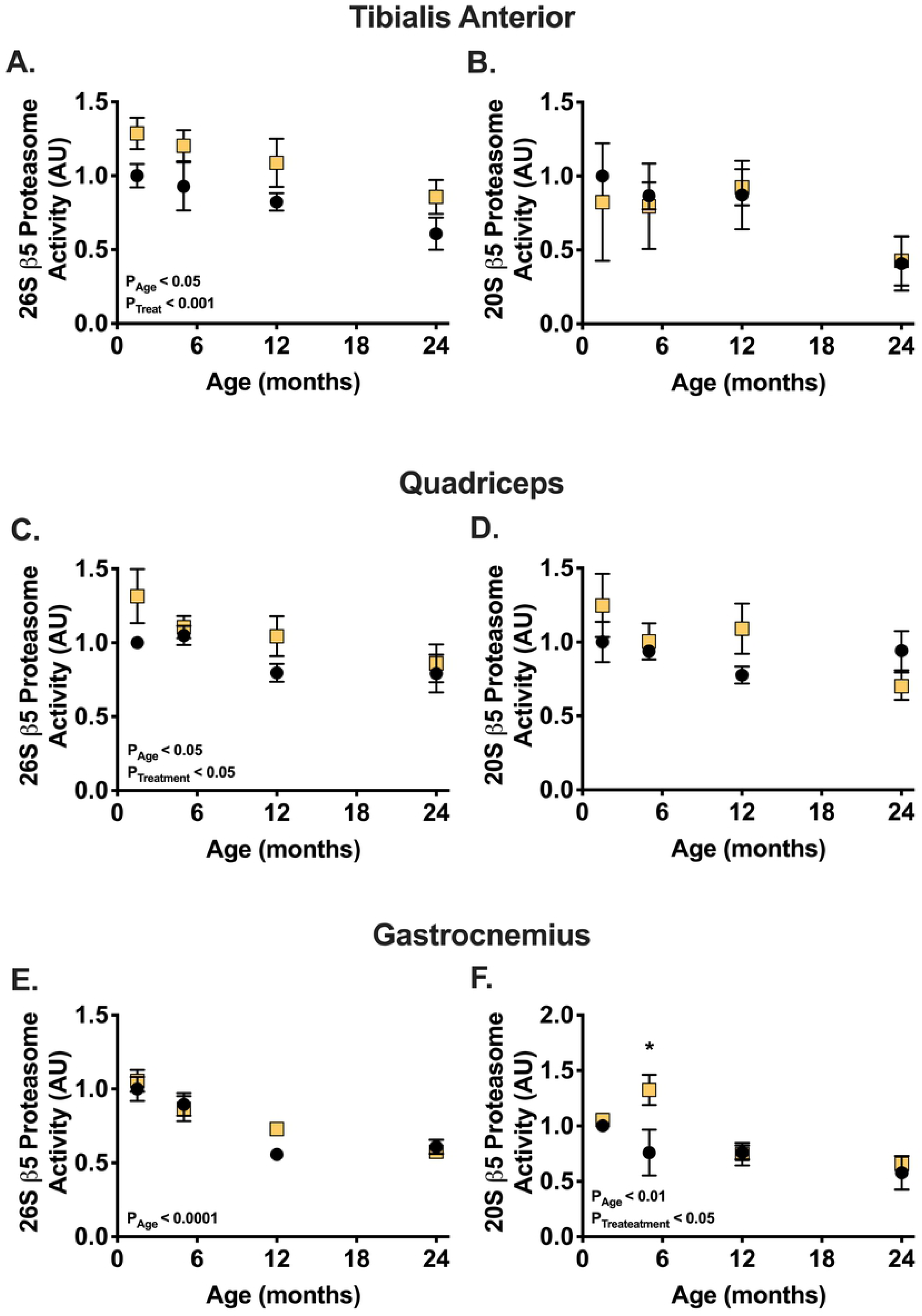
Protein degradation via the ubiquitin proteasome activity decreases with age and castration tends to increase the activity of this system. Proteasome activity of the 26S β-5 subunit in the tibialis anterior (A), quadriceps (C), and gastrocnemius (E), and. Proteasome activity of the 20S β-5 subunit in the tibialis anterior (B), and quadriceps (D), and gastrocnemius (F). * indicates p < 0.05 between treatment groups of the same age.

### Protein synthesis following one month of castration in male mice

Protein synthesis also plays an important role in determining protein balance. We therefore measured global rates of protein synthesis and estimated mTORC1 activity. The rate of protein synthesis, measured using the SUnSET method, was not affected in the three muscles following one month of testosterone depletion (Figures 5A, 5D, and 5G). Interestingly, the rate of protein synthesis increased in the gastrocnemius muscle of 5-month-old mice, regardless of testosterone levels, before decreasing back to a lower level (Figure 5G). 4E-BP1 phosphorylation by mTORC1 was also not affected by testosterone depletion (Figures 5B, 5E, 5H). Generally, starting at the age of 5-months, 4E-BP1 phosphorylation decreased with age in sham animals (Figures 5B, 5E, 5H).

**Fig 5.**
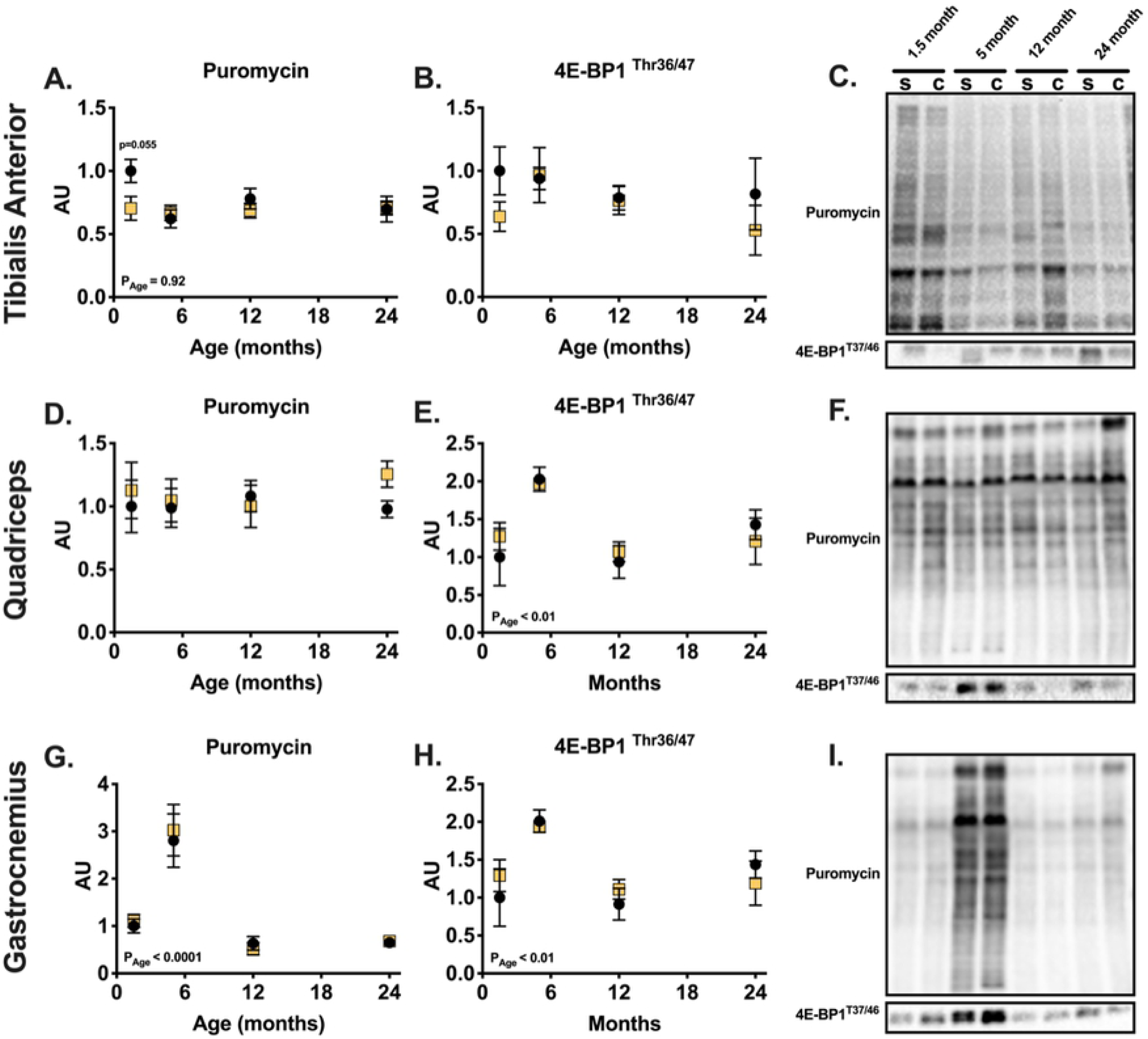
Basal levels of protein synthesis protein synthesis are not affected by testosterone depletion across life span and in different muscles. Quantification of protein synthesis rate using SUnSET in the tibialis anterior (A), quadriceps (D) and gastrocnemius (G) and quantification of 4E-BP1^The37/46^ phosphorylation in the tibialis anterior (B) quadriceps (E) and gastrocnemius (H) muscles. Representative SUnSET blots are shown to the right (C, F, I). * indicates p < 0.05 between treatment groups of the same age.

## Discussion

The present work supports our hypothesis that the effects of testosterone depletion on skeletal muscle mass change across the lifespan. These data show that testosterone is not important for the maintenance of skeletal muscle mass in adult male mice, but is critically important for early postnatal muscle growth. Testosterone depletion resulted in no change to muscle mass in 12- and 24-month-old animals. Moreover, sham animals exhibited a sarcopenic phenotype at 24 months in all muscles, a process that began at 12-months of age in the tibialis anterior muscle, with no effect of testosterone depletion. On the other hand, testosterone depletion halted natural growth in young juvenile male mice. Normal testosterone levels are crucial for early postnatal growth and development of skeletal muscle *in*-*vivo* (24). Our data demonstrate that testosterone serum concentrations rise in young mice, peaking around the age at which muscle mass peaks (Figures 1B, 1D, 1E, and 1F). This dynamic mirrors the trend in human males (24,25). It is therefore possible that the rise in testosterone serum concentrations supports musculoskeletal growth, a role that tappers-off following musculoskeletal maturity. This anabolic role of testosterone throughout postnatal growth may also be reflected in experiments conducted *in-vitro*, which routinely show positive effects of testosterone supplementation on myoblasts and myotubes (26). Cells used in *in*-*vitro* are in a developmental and high growth state, and therefore are more similar to skeletal muscle fibers in early postnatal animals (27,28).

Although testosterone may have a minimal role in the maintenance of muscle mass in adult animals (Figures 1D, 1E, and 1F), our data suggest that testosterone might be important in determining overall metabolism or the fate of adipose tissue (Figures 1A and 1C). Castration resulted in a significant increase in perirenal fat-pad mass at the same age. Although we do not have data available for other fat pads, we noticed a general increase in adiposity in 12-month-old sham animals compared to younger animals and this is reflected in their body weight (Figure 1A). As skeletal muscle mass was not higher in 12-month-old animals compared to 5-month-old animals, the increase in fat mass is likely the main contributor to the large increase in body weight we recorded between these groups. Moreover, both body weight and perirenal fat-mass significantly decreased in 12-month old animals following testosterone depletion. This suggests that following musculoskeletal maturity, testosterone may shift its role from an anabolic agent in skeletal muscle to regulator of fat accumulation and/or distribution. In fact, testosterone has been shown to effect adipose tissue in humans and in rats, with age being an important co-factor that modulates the effects (29–31). However, similar to its role in skeletal muscle, data regarding testosterone’s role in adipose tissue are inconclusive (31), and data from this study are limited and therefore precludes us from making a definitive statement regarding the role of testosterone in adipose tissue.

In contrast to the limited role testosterone plays in mature and aged skeletal muscle, this work clearly shows that low testosterone levels result in lower skeletal muscle mass in developing animals (Figures 1D, 1E, and 1F). The differences in muscle mass between sham and testosterone depleted animals may be due to either blunted growth or muscle mass loss in castrated animals. To address this question, we added a 4-month-old sham group. These animals are similar in size and muscle mass to the castrated 5-month cohort suggesting that testosterone depletion blunted growth, as opposed to inducing muscle atrophy. Although we do not have similar data for the younger cohort, we believe that testosterone depletion affected the 1.5-month-old mice similarly, essentially preventing further development. In past studies, testosterone depletion resulted in similar blunting of skeletal muscle growth in 2-month-old mice (18) and 3-6 month-old (32), but differences were interpreted to be related to atrophy. By including an additional control group, we suggest that the outcome of testosterone depletion was not atrophy. Interestingly, we detected changes in fiber cross sectional area only in the TA muscle of 1.5-month-old animals (Figures 2A, and 3A). This demonstrates the significant and critical role that testosterone has in early postnatal growth and development of skeletal muscle. Much of early postnatal growth is axial and testosterone has been shown to determine muscle fiber length in fish (33). Thus, testosterone may be important for axial growth in murine models as well. The greater effect on fiber CSA in 1.5-month-old animals following castration may be a result of testosterone playing a role in mediating axial lengthening of the tibialis anterior myofibers. The causative relationship between muscle length and fiber CSA were recently highlighted by Jorgenson and Hornberger (34). Additionally, testosterone administration is associated with increased addition of myoblasts (35,36), which is a key part of secondary fiber formation (37). Thus, it is possible that the decrease circulating testosterone levels in young animals results in dampened myoblasts addition and ultimately a decrease in fiber CSA. All in all, it is clear that age should be considered heavily in studies aimed at investigating the role testosterone plays in skeletal muscle growth and hypertrophy. The attention to age and adequate controls is crucial for interpretation of outcomes.

Testosterone depletion decreased muscle mass and fiber cross sectional area only in young animals, yet the treatment increased ubiquitin proteasome activity in all age groups. Specifically, testosterone depletion increased ATP dependent, 26S, ß5 subunit activity of the TA muscle (Figure 4A). This data supports a previous hypothesis suggesting that testosterone acts as a glucocorticoid receptor antagonist and therefore protects skeletal muscle from loss of mass (14– 16). However, because the activity of this system increased in all age groups, the probable antagonistic effect on the glucocorticoid receptor does not explain the phenotypic differences between young and old animals following castration (Figure 1); we detected no interaction between age and treatment in determining basal proteasome activity, which further suggests that the effect on basal activity of the proteasome system is not the cause of the different response to testosterone depletion between age groups. Moreover, neither resting protein synthesis, as measured by puromycin for 30 minutes prior to euthanasia, nor estimated mTORC1 activity indicate major differences in response to testosterone depletion across the mouse lifespan (Figure 5). This lack of difference also fails to explain the different effects of testosterone depletion throughout the mouse lifespan. Hence, the study does not support our second hypothesis which predicted that possible age differences would be a result of age specific differences in protein degradation following testosterone depletion.

The established paradigm regarding skeletal muscle mass suggests that muscle mass is determined by the closely regulated balance between myofibrillar protein synthesis and protein degradation (38). The outcomes we observed following testosterone depletion could not be explained by the measurements we made to assess baseline protein synthesis and protein degradation, as determined by ubiquitin proteasome activity. A clear limitation to our study is that measurements were taken after 30 days of testosterone depletion and it is possible that changes occurred earlier as has been shown previously (11). Additionally, our measurement of protein synthesis using puromycin has limitations in that it only detects proteins that are rapidly turning over. Measurement of protein synthesis over a longer time period using other methods such as the incorporation of a stable isotope (e.g. deuterated water) into the diet might be more informative (39). Finally, because our animals were fed at libitum and were not anabolically or catabolically stimulated prior to tissue collection, we may have missed peak differences in response between our young and old groups. This is a major limitation in our study design and methods, and future work should address this shortcoming.

Aside from supporting the important role of testosterone in early postnatal growth of muscle and the diminished role of testosterone in the maintenance of muscle mass in adult animals; this study highlights interesting differences between muscle groups in response to testosterone depletion. While growth of all muscles tested was impaired in testosterone-depleted young animals, the tibialis anterior was the most affected muscle (Figures 1D, and 2A) followed by the quadriceps muscle (Figure 1E), and the gastrocnemius muscle (Figure 1F). These data suggest that the effects of testosterone on skeletal muscle may be modulated by other factors such as external load or fiber type. In C57Bl6 mice, the tibialis anterior muscle receives less load during normal locomotion and has a greater proportion of the faster fiber types (IIb and IIx) compared to the gastrocnemius muscle (40). Because the tibialis anterior was more responsive to testosterone depletion compared to the gastrocnemius muscle, two possible hypotheses can be drawn. The first is that more oxidative muscles are less responsive to testosterone depletion. The second is that loaded muscles are less responsive to testosterone depletion. Another skeletal muscle that is routinely used to investigate the effects of testosterone supplementation and depletion on muscle mass, and is highly responsive to these treatments is the levator ani (41). The levator ani is not subjected to high loads and is a homogenously fast, type IIB muscle. The levator ani muscle has been shown to be more responsive to both testosterone depletion and supplementation compared to both the EDL (phenotypically faster), and soleus (phenotypically slower) (42). Additionally, previous data suggest that expression of the E3 ligases, MuRF-1 and MAFbx, was lower in loaded soleus and EDL muscles compared to the less loaded levator ani muscle in castrated c57Bl/6JOla mice that were supplemented with testosterone (12).Thus these data suggest that the effects of testosterone on adult muscle is not uniform and that factors such as muscle type and physiological loading need to be considered when evaluating the impact of testosterone depletion and supplementation.

### Conclusions

Testosterone plays no role in the maintenance of skeletal muscle mass in adult mice following musculoskeletal maturity. In contrast, testosterone is crucial for normal skeletal muscle growth in developing male mice. Additionally, the question regarding testosterone role in maintenance of muscle mass in very old mice is still unanswered and requires additional study. Normal testosterone function in male mice clearly involves an inhibition of the proteasome system. Yet, the effect on basal activity of this system may not be the cause for the difference in skeletal muscle phenotype following testosterone depletion between young and aging animals. Additionally, the role of testosterone in skeletal muscle growth is minimized with increased habitual load. Future studies should further investigate how testosterone depletion modulates proteostasis over long periods of time. In addition, the relationship between changes in loading activity of specific muscles should be further explored.

## Acknowledgments

We are grateful to University of California Davis Clinical and Transitional Sciences Center for the financial support of this work. We thank Dr Lucas Smith for the use of his microscope.

---

**supplemental Figure 1.**
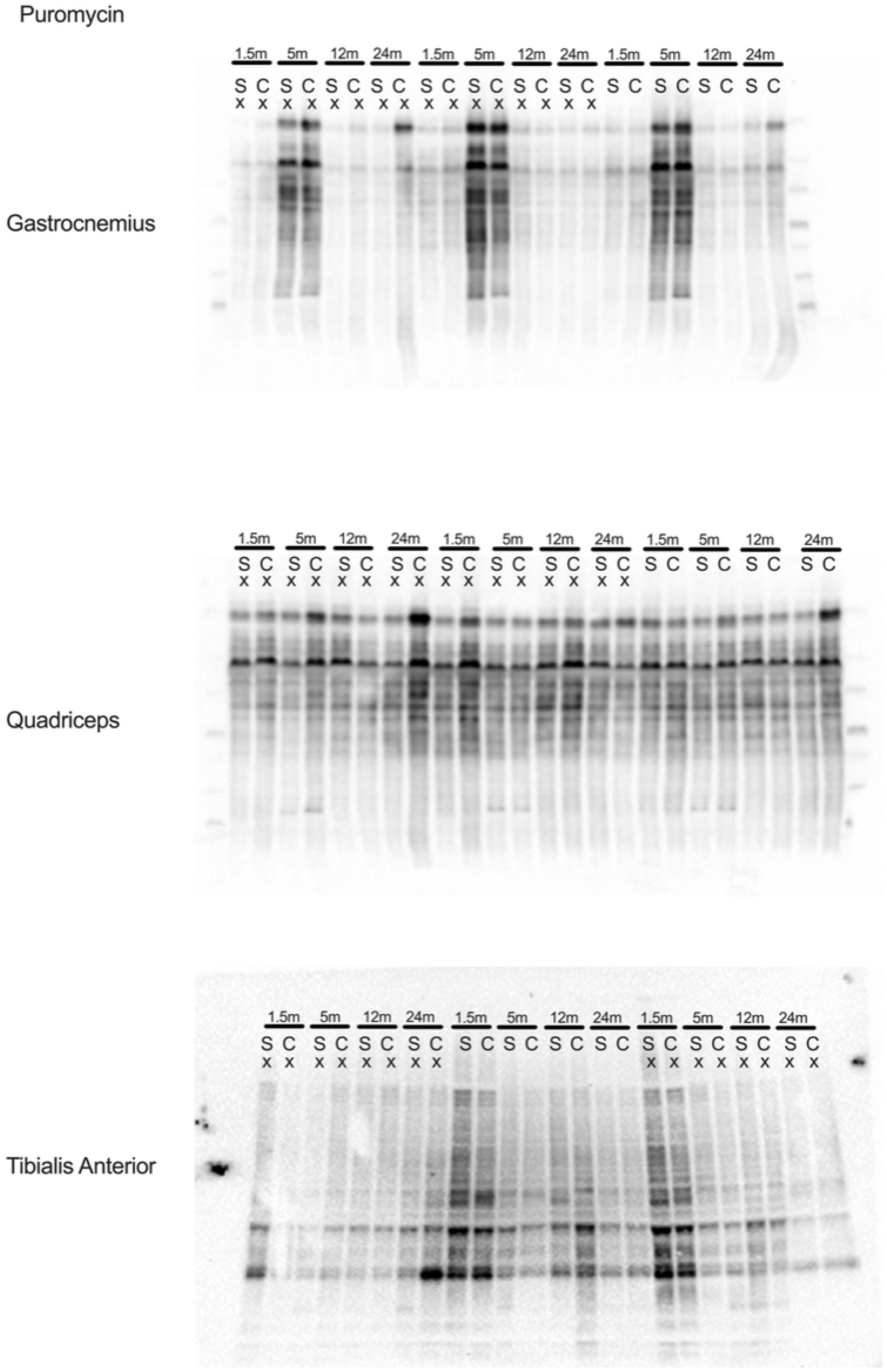

**supplemental Figure 2.**
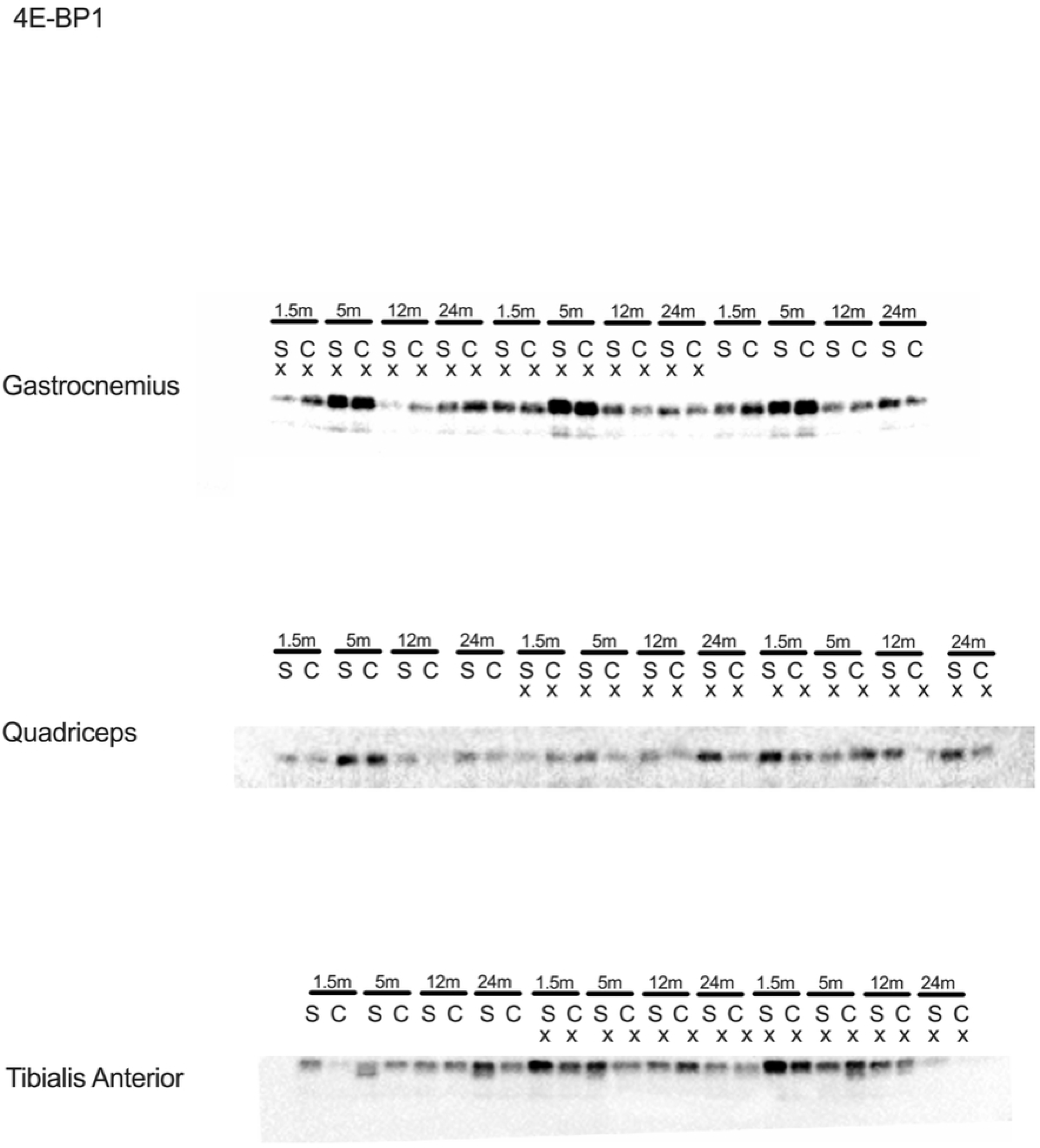

## References

1. Richter EA, Hargreaves M. Exercise, GLUT4, and Skeletal Muscle Glucose Uptake. Physiol Rev [Internet]. 2013;93(3):993–1017. Available from: http://physrev.physiology.org/cgi/doi/10.1152/physrev.00038.2012

2. Bonaldo P, Sandri M. Cellular and molecular mechanisms of muscle atrophy. Dis Model Mech [Internet]. 2013;6(1):25–39. Available from: http://www.pubmedcentral.nih.gov/articlerender.fcgi?artid=3529336&tool=pmcentrez&rendertype=abstract

3. Blaauw B, Schiaffino S, Reggiani C. Mechanisms modulating skeletal muscle phenotype. Compr Physiol. 2013;3(4):1645–87.

4. Clay C a, Perera S, Wagner JM, Miller ME, Nelson JB, Greenspan SL. Physical function in men with prostate cancer on androgen deprivation therapy. Phys Ther. 2007;87(10):1325–33.

5. Hohl A. Testosterone - From Basic to Clinical Aspects. Springer; 2017.

6. Shin MJ, Jeon YK, Kim IJ. Testosterone and Sarcopenia. World J Mens Health. 2018;36(3):192.

7. Herbst KL, Bhasin S. Testosterone action on skeletal muscle. Curr Opin Clin Nutr Metab Care. 2004;7(3):271–7.

8. Rossetti ML, Steiner JL, Gordon BS. Androgen-mediated regulation of skeletal muscle protein balance. Mol Cell Endocrinol [Internet]. 2017;447:35–44. Available from: http://dx.doi.org/10.1016/j.mce.2017.02.031

9. Kovacheva EL, Sinha Hikim AP, Shen R, Sinha I, Sinha-Hikim I. Testosteronesupplementation reverses sarcopenia in aging through regulation of myostatin, c-Jun NH2-terminal kinase, Notch, and Akt signaling pathways. Endocrinology. 2010;151(2):628–38.

10. White JP, Gao S, Puppa MJ, Sato S, Welle SL, Carson JA. Testosterone regulation of Akt/mTORC1/FoxO3a signaling in skeletal muscle. Mol Cell Endocrinol [Internet]. 2013;365(2):174–86. Available from: http://dx.doi.org/10.1016/j.mce.2012.10.019

11. Serra C, Sandor NL, Jang H, Lee D, Toraldo G, Guarneri T, et al. The effects of testosterone deprivation and supplementation on proteasomal and autophagy activity in the skeletal muscle of the male mouse: Differential effects on high-androgen responder and low-androgen responder muscle groups. Endocrinology. 2013;154(12):4594–606.

12. De Naeyer H, Lamon S, Russell AP, Everaert I, De Spaey A, Vanheel B, et al. Androgenic and estrogenic regulation of Atrogin-1, MuRF1 and myostatin expression in different muscle types of male mice. Eur J Appl Physiol. 2014;114(4):751–61.

13. Pires-Oliveira M, Maragno AL, Parreiras-e-Silva LT, Chiavegatti T, Gomes MD, Godinho RO. Testosterone represses ubiquitin ligases atrogin-1 and Murf-1 expression in an androgen-sensitive rat skeletal muscle in vivo. J Appl Physiol [Internet]. 2010;108(2):266–73. Available from: http://www.ncbi.nlm.nih.gov/pubmed/19926828%5Cnhttp://jap.physiology.org/content/jap/108/2/266.full.pdf

14. Zhao W, Pan J, Zhao Z, Wu Y, Bauman WA, Cardozo CP. Testosterone protects against dexamethasone-induced muscle atrophy, protein degradation and MAFbx upregulation. J Steroid Biochem Mol Biol. 2008;110(1–2):125–9.

15. Mayer M, Rosen F. Interaction of anabolic steroids with glucocorticoid receptor sites in rat muscle cytosol. Am J Physiol. 1975;229(5):1381–6.

16. Bodine SC, Furlow JD. Glucocorticoid Signaling [Internet]. Vol. 872, Advances in experimental medicine and biology. 2015. 253–78 p. Available from: http://www.ncbi.nlm.nih.gov/pubmed/26215998

17. Haren MT, Siddiqui AM, Armbrecht HJ, Kevorkian RT, Kim MJ, Haas MJ, et al. Testosterone modulates gene expression pathways regulating nutrient accumulation, glucose metabolism and protein turnover in mouse skeletal muscle. Int J Androl. 2011;34(1):55–68.

18. Rossetti ML, Steiner JL, Gordon BS. Increased mitochondrial turnover in the skeletal muscle of fasted, castrated mice is related to the magnitude of autophagy activation and muscle atrophy. Mol Cell Endocrinol [Internet]. 2018;3:1–8. Available from: https://doi.org/10.1016/j.mce.2018.01.017

19. Dalbo VJ, Roberts MD, Mobley CB, Ballmann C, Kephart WC, Fox CD, et al. Testosterone and trenbolone enanthate increase mature myostatin protein expression despite increasing skeletal muscle hypertrophy and satellite cell number in rodent muscle. Andrologia. 2017;49(3):1–11.

20. Jiao Q, Pruznak AM, Huber D, Vary TC, Lang CH. Castration differentially alters basal and leucine-stimulated tissue protein synthesis in skeletal muscle and adipose tissue. Am J Physiol Metab. 2009;297(5):E1222–32.

21. Dubois V, Laurent MR, Jardi F, Antonio L, Lemaire K, Goyvaerts L, et al. Androgen Deficiency Exacerbates High Fat Diet-Induced Metabolic Alterations in Male Mice. Endocrinology [Internet]. 2015;(November):en.2015–1713. Available from: http://press.endocrine.org/doi/10.1210/en.2015-1713

22. Goodman CA protein synthesis with SUnSET: a valid alternative to traditional techniques? Exerc Sport Sci Rev. 2013;41(2):107–15.

23. Gomes A V., Waddell DS, Siu R, Stein M, Dewey S, Furlow JD, et al. Upregulation of proteasome activity in muscle RING finger 1-null mice following denervation. FASEB J [Internet]. 2012;26(7):2986–99. Available from: http://www.fasebj.org/cgi/doi/10.1096/fj.12-204495

24. Liverman CT, Hamlin B EJ. Testosterone and Aging: Clinical Research Directions [Internet]. 2004. 240 p. Available from: https://books.google.com/books?id=BW2dAgAAQBAJ&pgis=1

25. Stárka L, Pospíšilová H, Hill M. Free testosterone and free dihydrotestosterone throughout the life span of men. J Steroid Biochem Mol Biol. 2009;116(1–2):118–20.

26. Basualto-Alarcón C, Jorquera G, Altamirano F, Jaimovich E, Estrada M. Testosterone signals through mTOR and androgen receptor to induce muscle hypertrophy. Med Sci Sports Exerc. 2013;45(9):1712–20.

27. Chal J, Pourquié O. Making muscle: Skeletal myogenesis in vivo and in vitro. Dev. 2017;144(12):2104–22.

28. Yaffe D, Ora S. Serial passaging of myogenic cells isolated from dystrophic mouse muscle. Nature [Internet]. 1977;270(December):11–3. Available from: https://www.nature.com/articles/270725a0.pdf

29. Kelly DM, Jones TH. Testosterone and obesity. Obes Rev [Internet]. 2015;(July):n/a-n/a. Available from: http://doi.wiley.com/10.1111/obr.12282

30. Traish AM, Miner MM, Morgentaler A, Zitzmann M. Testosterone deficiency. Am J Med [Internet]. 2011;124(7):578–87. Available from: http://dx.doi.org/10.1016/j.amjmed.2010.12.027

31. O’Reilly MW, House PJ, Tomlinson. JW. Understanding androgen action in adipose tissue. J Steroid Biochem Mol Biol [Internet]. 2014;143:277–84. Available from: http://dx.doi.org/10.1016/j.jsbmb.2014.04.008

32. Klose A, Liu W, Paris ND, Forman S, Krolewski JJ, Nastiuk KL, et al. Castration induces satellite cell activation that contributes to skeletal muscle maintenance. J Cachexia. 2018;1(1):1–13.

33. Akster HA. Numbers of Myosatellite Cells in White Axial Muscle of Growing Fish : Cyprinus carpio L. (Teleostei). 1991;424:418–24.

34. Jorgenson KW, Hornberger TA. The Overlooked Role of Fiber Length in Mechanical Load-Induced Growth of Skeletal Muscle. Exerc Sport Sci Rev [Internet]. 2019;47(4):258–9. Available from: http://www.ncbi.nlm.nih.gov/pubmed/31524787%0Ahttp://www.pubmedcentral.nih.gov/articlerender.fcgi?artid=PMC6750012

35. Sinha-Hikim I, Artaza J, Woodhouse L, Gonzalez-Cadavid N, Singh AB, Lee MI, et al. Testosterone-induced increase in muscle size in healthy young men is associated with muscle fiber hypertrophy. Am J Physiol Endocrinol Metab. 2002;283(1):E154–64.

36. Sinha-Hikim I, Roth SM, Lee MI, Bhasin S. Testosterone-induced muscle hypertrophy is associated with an increase in satellite cell number in healthy, young men. Am J Physiol - Endocrinol Metab [Internet]. 2003;285(148-1):197–205. Available from: http://ajpendo.physiology.org/lookup/doi/10.1152/ajpendo.00370.2002

37. Fredette BJ, Landmesser LT. Relationship of primary and secondary myogenesis to fiber type development in embryonic chick muscle. Dev Biol. 1991;143(1):1–18.

38. Sandri M. Signaling in Muscle Atrophy and Hypertrophy. Physiology [Internet]. 2008;23(3):160–70. Available from: http://physiologyonline.physiology.org/cgi/doi/10.1152/physiol.00041.2007

39. Wilkinson DJ, Brook MS, Smith K, Atherton PJ. Stable isotope tracers and exercise physiology: Past, present and future. J Physiol [Internet]. 2016;00(March):1–10. Available from: http://www.ncbi.nlm.nih.gov/pubmed/27610950

40. Augusto V, Padovani CR, Eduardo G, Campos R. SKELETAL MUSCLE FIBER TYPES IN C57BL6J MICE. 2004;21:89–94.

41. Gori Z, Pellegrino C, Pollera M. The hypertrophy of levator ani muscle of rat induced by testosterone: an electron microscope study. Exp Mol Pathol [Internet]. 1969;10(2):199–218. Available from: http://www.ncbi.nlm.nih.gov/entrez/query.fcgi?cmd=Retrieve&db=PubMed&dopt=Citation&list_uids=5777797

42. Axell A-M, MacLean HE, Plant DR, Harcourt LJ, Davis J a, Jimenez M, et al. Continuous testosterone administration prevents skeletal muscle atrophy and enhances resistance to fatigue in orchidectomized male mice. Am J Physiol Endocrinol Metab [Internet]. 2006;291(3):E506–16. Available from: http://www.ncbi.nlm.nih.gov/pubmed/16621900

